# Midbrain neurons regulate the production of courtship squeaks in female mice

**DOI:** 10.64898/2026.11.06.764661

**Authors:** Da-Jiang Zheng, Renee J. Henderson, Patryk Ziobro, Zizhan Zhou, Katherine A. Tschida

**Affiliations:** Department of Psychology, Cornell University, 109 Tower Rd., Ithaca NY 14853

**Keywords:** vocalization, female, courtship, squeak, periaqueductal gray

## Abstract

Animals produce different vocalization types (e.g., contact calls, mating calls) to communicate in different behavioral contexts, but how the brain is organized to enable the production of different vocalization types remains poorly understood. Recent work found that specialized neurons in the midbrain periaqueductal gray (PAG) regulate the production of ultrasonic vocalizations (USVs) in female and male mice, but these PAG-USV neurons are not required for the production of broadband vocalizations (squeaks). To identify PAG neurons that may regulate the production of squeaks, we used an activity-dependent labeling strategy (TRAP2) to identify and manipulate PAG neurons that are active in association with the production of courtship squeaks in female mice. Optogenetic activation of courtship squeak-TRAPed PAG neurons elicits the production of squeaks, while ablation of TRAPed PAG neurons blocks the production of courtship squeaks without altering female non-vocal courtship behaviors. Surprisingly, ablation of courtship squeak-TRAPed PAG neurons did not affect the production of footshock-elicited squeaks. Moreover, non-selective PAG ablation, which blocked the production of courtship squeaks in females and USVs in both sexes, attenuated but did not block the production of footshock-elicited squeaks. Our results identify a population of PAG neurons that critically regulates the production of courtship squeaks in female mice and also suggest that a non-canonical, PAG-independent pathway regulates the production of pain-elicited vocalizations.

## Introduction

Vocalizations are important for communication in both human and non-human animals. To communicate flexibly across different behavioral contexts, animals produce a variety of vocalization types, which differ in their acoustic features (e.g., contact calls when separated from conspecifics, mating calls during courtship interactions, alarm calls when a predator is detected). How the brain is organized to enable the production of different vocalization types remains poorly understood.

The midbrain periaqueductal gray (PAG) is a key brain region that regulates the production of vocalizations in vertebrates, including in mammals, birds, and fish.^1–7^ Artificial activation of the PAG elicits the production of species-typical vocalizations, and activation at different PAG sites can in some cases elicit the production of different vocalization types.^1,8,9^ On the other hand, silencing of the PAG blocks the production of species-typical vocalizations, and silencing at different PAG sites can block the production of certain vocalization types while leaving others intact.^1,10–12^ These studies suggest that at least some vocalization types may depend on different populations of PAG neurons, but resolving this issue requires the identification of specific PAG neurons that are important for a given vocalization type in order to test their functional role in regulating additional vocalization types.

Recent work in mice has begun to address this question. Adult mice produce two main vocalization types. First, ultrasonic vocalizations (USVs) are produced during social interactions, particular by males when they engage in courtship interactions with females, as well as by females during same-sex interactions.^13–18^ Second, mice produce lower-frequency, broadband vocalizations (i.e., squeaks) when in pain and/or during interactions with aggressive conspecifics or predators,^19–23^ and female mice also produce squeaks during courtship interactions with males.^21,24^ A pair of studies showed that specialized PAG neurons regulate the production of USVs. Activation of these PAG-USV neurons elicits USV production, while silencing or ablation of PAG-USV neurons blocks the production of USVs in both female and males.^23,25^ Although PAG-USV neurons are required for USV production, caspase-mediated ablation of PAG-USV neurons does not block the production of squeaks, either those elicited by mild footshock in females and males or courtship squeaks produced by females.^23^ Given that the PAG is thought to regulate the production of all innate vocalizations, these results suggest that a distinct population of PAG neurons regulates the production of squeaks. However, such neurons remain to be identified.

In the current study, we used an intersectional approach that enables the expression of viral transgenes in recently active neurons (TRAP2)^26,27^ to identify PAG neurons that regulate squeak production in mice. One challenge to using TRAP2 to identify such neurons is that in addition to regulation vocalization, the PAG regulates nociception,^28,29^ and most squeak-eliciting behavioral contexts involve pain (e.g., experimenter-delivered footshock, being bitten by an aggressive conspecific). Moreover, efficient TRAPing of any squeak-related PAG neurons likely requires high rates of squeak production, but eliciting high squeak rates in pain-associated contexts risks injury to the subject animals. To circumvent these challenges, we applied the TRAP2 strategy in female mice that produced high rates of courtship squeaks during interactions with sexually motivated males.^30^ Although we leveraged the courtship squeak context to identify squeak-related PAG neurons, we hypothesized that the identified neurons would regulate the production of all squeaks, even those produced outside of a courtship context.

## Results

### Caudolateral PAG neurons upregulate Fos expression following the production of courtship squeaks in female mice

To localize PAG neurons that may regulate the production of squeaks, we first determined whether the PAG contains neurons that upregulate expression of the immediate early gene Fos in association with the production of female courtship squeaks. Female subject mice were given courtship interactions with sexually motivated male mice (Fig. 1A; see Methods), a context in which female mice naturally produce high rates of squeaks, predominantly in association with rejection of male mounting attempts.^21,24,31^ We quantified Fos expression across the rostral-to-caudal extent of lateral and dorsal PAG, because the lateral PAG contains neurons that project to the nucleus retroambiguus,^25,32^ a hindbrain region that is required for the production of squeaks.^32^ Dorsolateral PAG has been implicated in regulating squeak production in high anxiety mice,^22^ and dorsomedial PAG regulates female rejection behaviors during courtship interactions with males.^33^ We compared PAG Fos expression (lateral and dorsal PAG pooled initially) in females that were given brief (<5 minutes) interactions with males and produced modest rates of courtship squeaks (n = 6 low squeak females; mean = 35 squeaks, range = 7-73 squeaks) and in females that were given long (30-minute) interactions with males and produced high rates of courtship squeaks (n = 9 high squeak females; mean = 200 squeaks, range = 82-441 squeaks) (Fig. 1B-C). This comparison revealed that high squeak females tended to have higher Fos expression throughout the rostral-to-caudal extent of the PAG, although this difference overall was not significant (Fig. 1D; Mann Whitney U test, total PAG Fos counts in low vs. high squeak females; z = 1.12, p = 0.13). We noted, however, that the trend toward a difference in total Fos-positive PAG neurons between the two groups seemed more pronounced in the caudal half than in the rostral half of the PAG. We therefore repeated the analysis separately for rostral PAG (–3.4 to –4.0 mm A/P) and for caudal PAG (–4.2 to –5.0 mm A/P). In rostral PAG, we found that the total number of Fos-positive neurons did not differ between the two groups (Fig. 1E, left; Mann Whitney U test; z = 0.65, p = 0.26), nor was the total number of Fos-positive neurons significantly related to total squeaks produced by each female (Fig. 1E, right; linear regression after log-transforming squeak counts; t(13) = 1.57, p = 0.14). In contrast, the total number of Fos-positive neurons in the caudal PAG differed significantly between the two groups (Fig. 1F, left; Mann Whitney U test, z = 1.71, p = 0.04), and the total number of Fos-positive neurons was significantly related to total squeaks produced by each female (Fig. 1F, right; linear regression after log-transforming squeak counts; t(13) = 2.94, p = 0.01). Moreover, the relationship between caudal PAG Fos expression and total squeaks produced by each female was stronger for caudolateral PAG neurons than for caudodorsal PAG neurons (linear regressions after log-transforming squeaks counts: t(13) = 3.49, p = 0.004 for caudolateral PAG; t(13) = 1.76, p = 0.10 for caudodorsal PAG). We conclude that caudolateral PAG neurons upregulate Fos expression in association with the production of female courtship squeaks.

**Figure 1.**
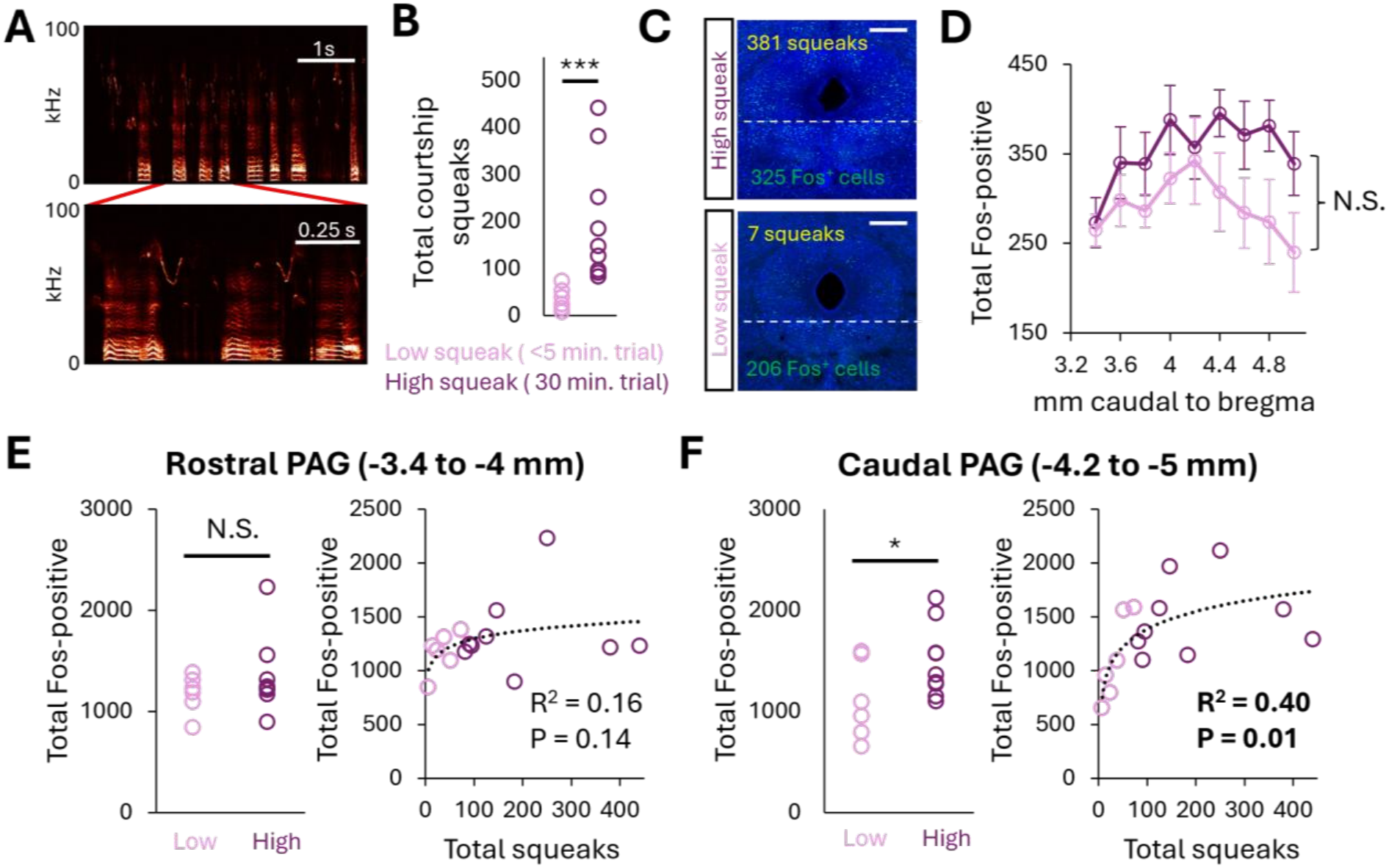
Fos expression in the caudal PAG is correlated with the production of female courtship squeaks. (A) Spectrograms show example female courtship squeaks. (B) Plot shows total courtship squeaks produced by females given < 5-minute-long interactions with males (n = 6 low squeak) and females given 30-minute-long interactions with males (n = 9 high squeak). (C) Confocal images show PAG Fos expression in example high squeak (top) and low squeak (bottom) females. Blue, neurotrace; green, Fos. Scale bars, 500 µm. Dotted lines indicate the ventral boundary for PAG Fos quantification. (D) Plot shows mean counts of Fos-positive PAG neurons across planes of coronal section, in low squeak and high squeak females. (E) Left plot shows summed counts of Fos-positive rostral PAG neurons for high squeak and low squeak females. Right plot shows relationship between total courtship squeaks and total Fos-positive rostral PAG neurons across females. (F) Same as E, for Fos-positive caudal PAG neurons.

### Optogenetic activation of courtship squeak-TRAPed PAG neurons robustly elicits the production of squeaks in female mice

Are these Fos-positive caudolateral PAG neurons functionally relevant to the production of squeaks? To test whether artificial activation of Fos-positive PAG neurons is sufficient to elicit the production of squeaks, we used the TRAP2 activity-dependent labeling strategy to express channelrhodopsin (ChR2) in PAG neurons that upregulate Fos in association with the production of female courtship squeaks. Briefly, female TRAP2 mice received unilateral PAG injections of AAV-FLEX-ChR2 and were implanted with a fiber optic over the lateral PAG (Fig. 2A-B). Two weeks later, female subjects were given a 30-minute courtship interaction with a sexually motivated male, produced robust rates of courtship squeaks (Fig. 2C), and then received an IP injection of 4-hydroxytamoxifen (4-OHT), which enables the transient expression of Cre recombinase in recently active neurons and therefore permits the expression of ChR2 in courtship squeak-TRAPed PAG neurons. Two weeks later, we then tested whether optogenetic activation of courtship squeak-TRAPed PAG neurons was capable of eliciting the production of squeaks in solo-tested females. Control females were treated equivalently, except that they received unilateral PAG injections of AAV-FLEX-GFP.

**Figure 2.**
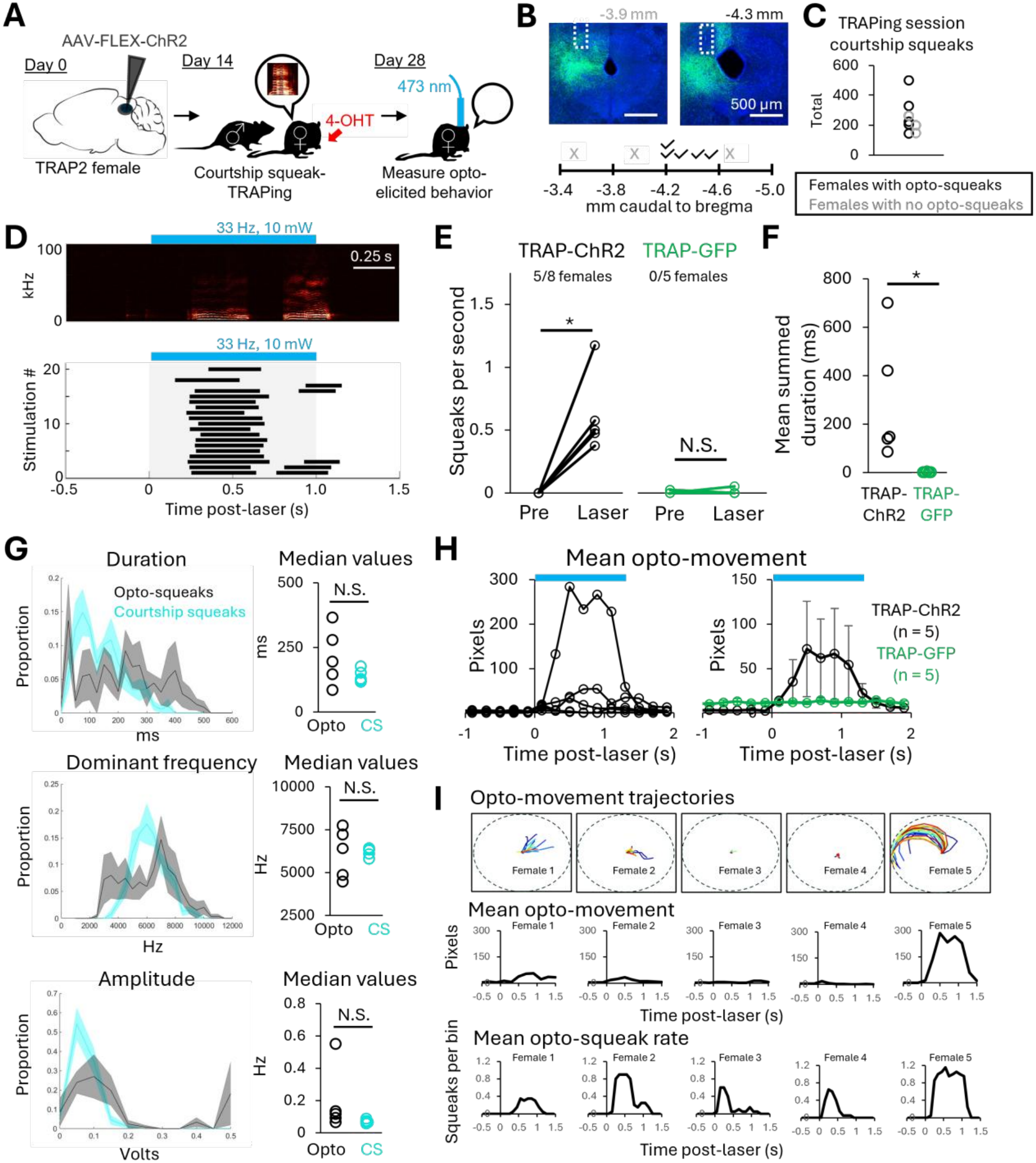
Optogenetic activation of courtship-squeak-TRAPed PAG neurons elicits the production of squeaks in female mice. (A) Schematic shows experimental details and time course. (B) Confocal images show viral labeling and fiber optic placement in a TRAP-ChR2 female that did not exhibit opto-elicited squeaks (left) and for a TRAP-ChR2 female that exhibited opto-elicited squeaks (right). Plot below shows ferrule placement (mm caudal to bregma) for TRAP-ChR2 females that exhibited opto-elicited squeaks (n = 5, black symbols) and for TRAP-ChR2 females that did not exhibit opto-elicited squeaks (n = 3, gray symbols). Scale bars, 500 µm. (C) Plot shows total number of courtship squeaks produced during TRAPing sessions for TRAP-ChR2 females that subsequently exhibited opto-elicited squeaks (black symbols) and for TRAP-ChR2 females that did not exhibit opto-elicited squeaks (gray symbols). (D) Spectrogram shows representative squeaks elicited by optogenetic activation of courtship squeak-TRAPed PAG neurons. Raster plot below shows timing of opto-elicited squeaks in all trials from the same TRAP-ChR2 female. Blue bars show timing of laser. (E) Plot shows squeaks per second produced during pre-laser baseline periods (2s) and during laser periods (includes 1s laser stimulation and 1s post-stimulation) for TRAP-ChR2 experimental females and TRAP-GFP control females. (F) Plots show mean summed duration of optogenetically-elicited squeaks produced by TRAP-ChR2 and TRAP-GFP females. (G) (Left) Distributions of three acoustic parameters are shown for optogenetically-elicited squeaks produced by TRAP-ChR2 females (black traces) and for courtship squeaks produced by unmanipulated females recorded in the same behavioral chamber (cyan traces). (Right) Plots show median values of the same three acoustic parameters for opto-elicited squeaks produced by TRAP-ChR2 females and for courtship squeaks produced by unmanipulated females. (H) (Left) Plot shows mean movement elicited by optogenetic stimulation of courtship-TRAPed PAG neurons for each TRAP-ChR2 female. (Right) Plot shows group means of movement elicited by optogenetic stimulation of courtship-TRAPed PAG neurons for TRAP-ChR2 females and for TRAP-GFP females. (I) Top row shows trajectories of opto-elicited movements for each TRAP-ChR2 female, from laser onset to laser offset. Middle row shows plots of mean opto-elicited movement for each TRAP-ChR2 female. Bottom row shows plots of opto-elicited squeak rate for each TRAP-ChR2 female.

In the majority (5 of 8) of TRAP-ChR2 females, we observed that optogenetic activation of courtship squeak-TRAPed PAG neurons elicited the production of loud, robust squeaks (Fig. 2D-E, Video S1; 1s-long laser stimulations, 33 Hz, 10 mW). When we examined the targeting of PAG viral injections and fiber optic placements across these cases, we found that the 5 females that exhibited optogenetically-elicited squeaks were injected/implanted between –4.0 and –4.5 mm A/P, while the 3 females that failed to exhibit optogenetically-elicited squeaks had targeting that was either rostral to (n = 2) or caudal to this range (n = 1) (Fig. 2B). All TRAP-ChR2 females produced robust rates of courtship squeaks during their TRAPing sessions (Fig. 2C), indicating that differences in TRAPing session vocal behavior do not account for the variability in experimental outcomes. In contrast to the robust squeaks elicited by laser stimulation in the n = 5 TRAP-ChR2 females, laser stimulation failed to elicit squeaks in TRAP-GFP control females (Fig. 2E-F; Mann Whitney U test on the difference between laser squeak rates and baseline squeak rates for each group; z = 2.51, p = 0.01; Mann Whitney U test on mean summed squeak durations; z = 2.51, p = 0.01). Optogenetically-elicited squeaks did not differ significantly from natural courtship squeaks produced by unmanipulated control females in their median duration, dominant frequency, or amplitude (Mann Whitney U tests; p > 0.05 for all acoustic features), but histogram plots of these acoustic features highlight that optogenetically-elicited squeaks are overall more variable in their acoustic features than naturally-produced courtship squeaks (Fig. 2G; F-tests for equality of variance; p < 0.05 for all acoustic features). In summary, optogenetic activation of courtship squeak-TRAPed PAG neurons is capable of eliciting squeak production in female mice.

Mice often produce squeaks in association with active defensive behaviors (e.g., fleeing from an aggressive conspecific or predator, jumping in response to footshock) but not during passive defensive behaviors (e.g., freezing). Moreover, females exhibit a number of active defensive behaviors during courtship interactions with males (fleeing, boxing, kicking, etc.). We therefore wondered whether optogenetically-elicited squeaks produced by the TRAP-ChR2 females were coupled to or produced in a manner that was secondary to active defensive behaviors. In addition to eliciting the production of squeaks, we found that optogenetic activation of courtship squeak-TRAPed PAG neurons elicited body movements, but in a highly variable manner across females. One of the TRAP-ChR2 females exhibited large amplitude, escape-like locomotion upon optogenetic stimulation (female 5), whereas other females exhibited small body movements (female 1), small head movements and infrequent, small body movements (female 2), or freezing (females 3 and 4) (Fig. 2H-I; see Fig. S1 for visualizations of trajectories of optogenetically-elicited movements in all TRAP-ChR2 and TRAP-GFP females). TRAP-ChR2 females also exhibited freezing in between optogenetic stimulations, which can be seen by comparing the pre-laser movement between the TRAP-ChR2 experimental females and the TRAP-GFP control females (Fig. 2H, right). Although these data indicate that optogenetic activation of courtship squeak-TRAPed PAG neurons does not exclusively elicit squeak production, the variable effects on movement alongside the consistent and robust effects on squeak production support the idea that the production of optogenetically-elicited squeaks is not coupled to or secondary to the production of active defensive behaviors.

We also tested the effects of optogenetic activation of courtship squeak-TRAPed PAG neurons during interactions with males, to test whether neuronal activation would result in different or additional behaviors than those observed in solo optogenetic sessions. In these trials, we used males that were sexually inexperienced, to ensure that any squeaks produced by the female were optogenetically-elicited. These males engaged in social investigation of subject females but did not make mounting attempts. Accordingly, all squeaks were produced by the females during periods of laser stimulation. Squeak rates did not differ significantly between solo and male partner optogenetic sessions (Fig. S2A; paired t-test; t(4) = 2.19, p = 0.09), and the total amount of movement elicited by optogenetic stimulation did not differ between session types (Fig. S2B; t(4) = –0.09, p = 0.93). Moreover, the trajectories of optogenetically-elicited movements looked similar between session types (Fig. S2C), indicating that movement trajectories weren’t altered in a manner that depended on the position of the male relative to the female at the time of stimulation onset. We also did not observe the production of male-directed rejection behaviors, such as kicking or slapping (Video S2). We conclude that optogenetic activation of courtship squeak-TRAPed PAG neurons elicits squeak production and variable effects on body movement but does not elicit male-directed defensive behaviors.

### Courtship TRAPed-PAG neurons project to downstream regions implicated in vocal production

We next traced the axonal projections of courtship squeak-TRAPed PAG neurons (from n = 5 TRAP-ChR2 females that exhibited opto-elicited squeaks), to begin to understand how they engage with downstream circuits to promote squeak production. We found that the axons of these neurons target regions including the pontine reticular formation, the lateral parabrachial nucleus, the pontine tegmentum adjacent to locus coeruleus (peri-LC region), the gigantocellular reticular nucleus (alpha part and ventral part), the lateral paragigantocellular nucleus, and nucleus retroambiguus (RAm) (Fig. 3). Of these regions, both the peri-LC region^34^ and RAm^25,30,32,35–37^ have been implicated in regulating rodent USV production. Although the hindbrain neuronal populations that regulate squeak production remain less explored, recent studies have also implicated RAm in regulating the production of squeaks^30,32,37^. In summary, courtship squeak TRAPed PAG neurons send axonal projections to hindbrain areas implicated in regulating vocal production, including the production of squeaks.

**Figure 3.**
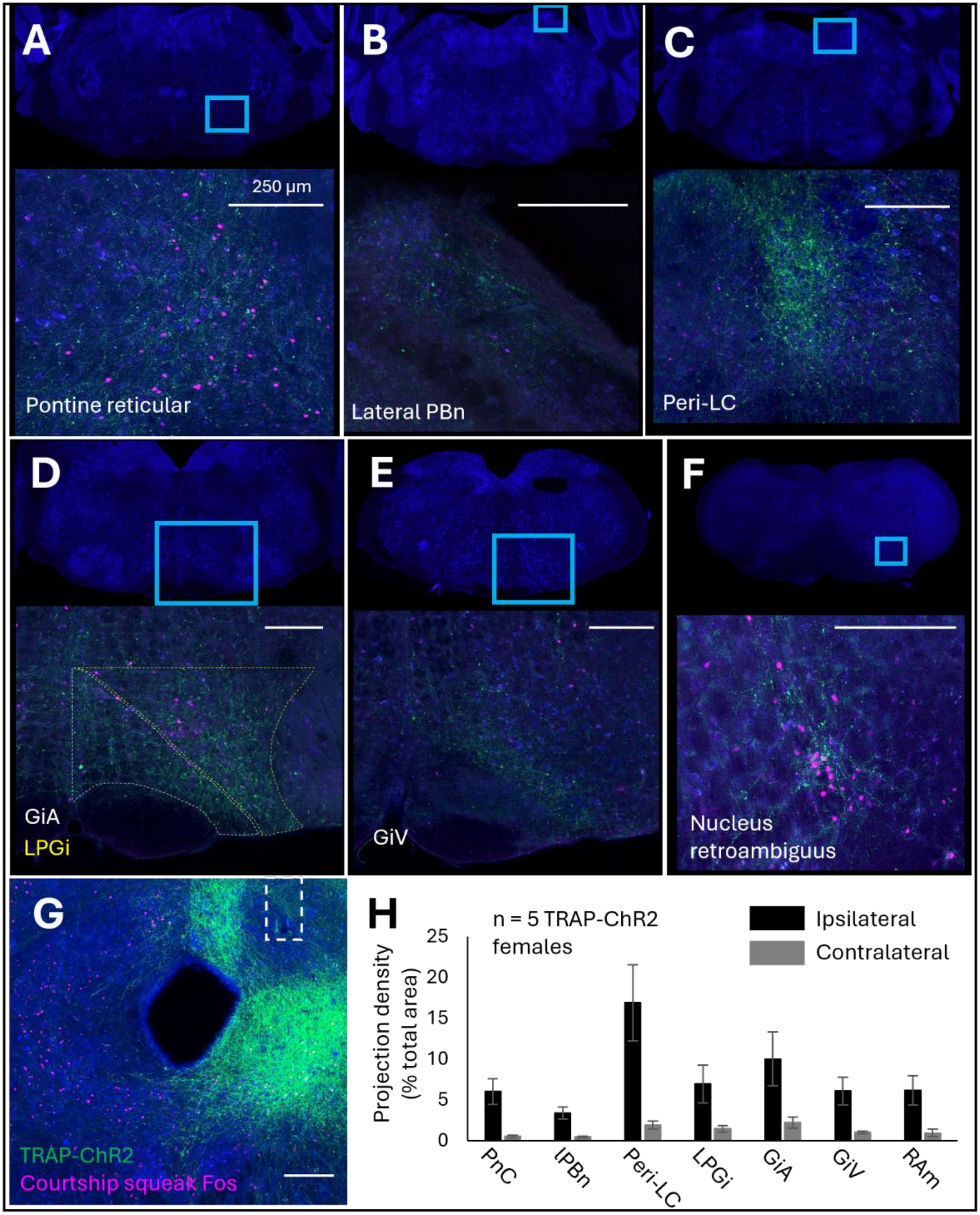
Axonal projections of courtship-squeak TRAPed PAG neurons. Representative confocal images show axonal projections to (A) the pontine reticular formation (PnC), (B) the lateral parabrachial nucleus (lPBn), the peri-locus coeruleus region (peri-LC), (D) the lateral paragigantocellular nucleus (LPGi; yellow dotted line) and the alpha part of the gigantocellular reticular nucleus (GiA; white dotted line), (E) the ventral part of the gigantocellular reticular nucleus (GiV), and nucleus retroambiguus (RAm). (G) Confocal image shows a representative PAG injection site. Dotted rectangle shows approximate placement of optical fiber. Green, axons of courtship TRAPed PAG neurons (labeled with AAV-FLEX-ChR2-EYFP; magenta, courtship squeak Fos. (H) Quantification of axonal projections, presented as projection density (total square pixels of axonal innervation normalized by the total area of each region, organized from rostral to caudal; mean ± SE). Scale bars, 250 µm.

### Optogenetic activation of VGlut2^+^ PAG neurons weakly elicits the production of squeaks in both female and male mice

Although our TRAP2-based optogenetic experiments were conducted exclusively in females, we wondered optogenetic activation of the same PAG region in which TRAP-based activation elicited squeaks (–4 to –4.5 mm A/P) using a non-TRAP-based strategy is capable of eliciting the production of squeaks in both sexes. To this end, we tested whether optogenetic activation of glutamatergic PAG neurons (VGlut2^+^, the predominant midbrain VGlut isoform^38^) was capable of eliciting squeak production in females and males (Fig. 4A). In contrast to the robust squeaks elicited by activation of courtship squeak-TRAPed PAG neurons, optogenetic activation of VGlut2^+^ PAG neurons elicited the production of squeaks in only 50% of the mice, but with equal likelihood for both sexes (Fig. 4B-C; optogenetically-elicited squeaks were observed in 4 of 8 VGlut2-ChR2 females, 3 of 6 VGlut2-ChR2 males, and 0 of 9 control VGlut2-GFP mice; Mann Whitney U test on difference between laser squeak rates and baseline squeaks rates for each group, z = –3.28, p = 0.001; laser stimulations were 3s-long, 33 Hz, 5 mW). Compared to TRAP-ChR2 females, VGlut2-ChR2 mice had a lower proportion of laser stimulations that elicited squeaks, had significantly lower rates of optogenetically-elicited squeaks (excluding laser stimulations with 0 opto-squeaks), and exhibited significantly longer latencies from laser onset to squeak (Fig. 4D; Mann Whitney U tests; p < 0.05 for all). Moreover, the optogenetically-elicited squeaks produced by VGlut2-ChR2 mice were significantly lower in both duration and amplitude than optogenetically-elicited squeaks produced by TRAP-ChR2 females (Fig. 4E; Mann Whitney U tests; p < 0.05 for both). We also noted that 1 of the 7 VGlut2-ChR2 mice that exhibited optogenetically-elicited squeaks also produced optogenetically-elicited USVs, but the low likelihood of obtaining USVs is likely because the targeting for the VGlut2-ChR2 mice is still rostral to the caudal-most portion of the PAG known to contain PAG-USV neurons^25,30,34,39^

**Figure 4.**
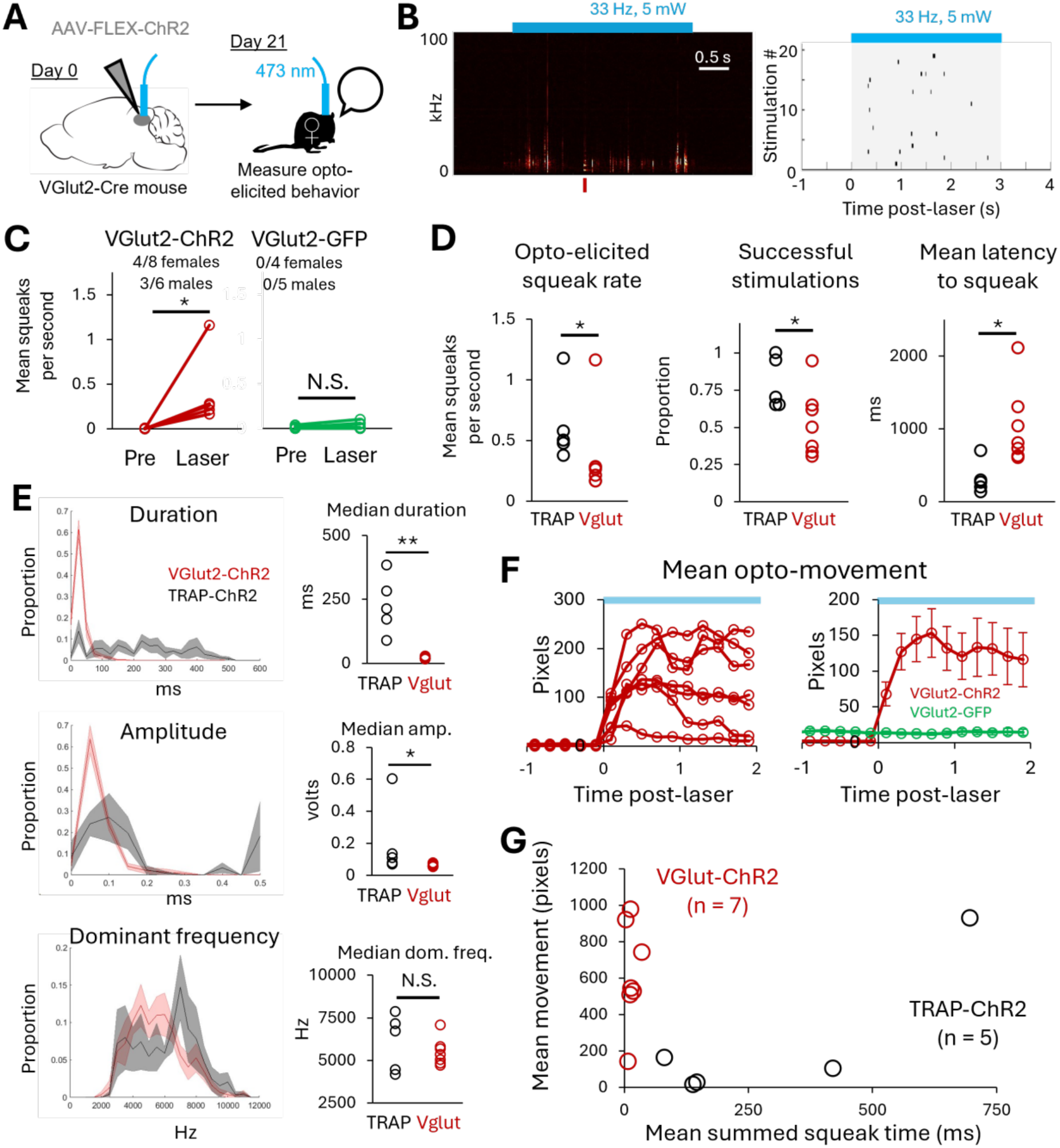
Optogenetic activation of VGlut2^+^ PAG neurons weakly elicits the production of squeaks in both females and males. (A) Schematic shows experimental details and time course. (B) Left, spectrogram shows representative squeaks elicited by optogenetic activation of VGlut2^+^ PAG neurons. Right, raster plot below shows timing of opto-elicited squeaks in all trials from the same VGlut2-ChR2 mouse. Blue bars show timing of laser. (C) Plot shows squeaks per second produced during pre-laser baseline periods (4s) and during laser periods (includes 3s laser stimulation and 1s post-stimulation) for VGlut2-ChR2 experimental mice (red symbols) and VGlut2-GFP control mice (green symbols). (D) Plots show mean rate of optogenetically-elicited squeaks (left), mean proportion of successful laser stimulations (middle), and mean latency from laser stimulation to first squeak (right) for VGlut2-ChR2 mice (red symbols) vs. TRAP-ChR2 females (black symbols). (E) (Left) Distributions of three acoustic parameters are shown for optogenetically-elicited squeaks produced by VGlut2-ChR2 mice (red traces) and by TRAP-ChR2 females (black traces). (Right) Plots show median values of the same three acoustic parameters for optogenetically-elicited squeaks produced by VGlut2-ChR2 mice and by TRAP-ChR2 females. (F) (Left) Plot shows mean movement elicited by optogenetic stimulation of Vglut2^+^ PAG neurons for each VGlut2-ChR2 mouse that exhibited optogenetically-elicited squeaks. (Right) Plot shows group mean movement elicited by optogenetic stimulation of VGlut2^+^ PAG neurons for VGlut2-ChR2 mice and for VGlut2-GFP mice. (G) Plots show mean summed squeak time vs. mean movement for each VGlut2-ChR2 mouse and for each TRAP-ChR2 female. Because VGlut2-ChR2 received longer laser stimulations than TRAP-ChR2 (3s vs. 1s), this analysis includes only the first second of behavior after laser onset for each mouse.

In contrast to relatively weak effects on squeak production, optogenetic activation of VGlut2^+^ PAG neurons elicited robust non-vocal movement, with each mouse exhibiting large amplitude, escape-like locomotion upon optogenetic stimulation (Fig. 4F, Video S3; see Fig. S1 for visualizations of trajectories of optogenetically-elicited movements). In summary, optogenetic activation of VGlut2^+^ PAG weakly elicits squeaks and robustly elicits non-vocal movements in both females and males, whereas optogenetic activation of courtship squeak-TRAPed PAG neurons robustly elicits squeaks and exerts variable and relatively weaker effects on non-vocal movements (Fig. 4G).

### Ablation of courtship squeak-TRAPed PAG neurons blocks the production of courtship squeaks, but not footshock-elicited squeaks, in female mice

We next tested whether the activity of courtship squeak-TRAPed PAG neurons is required for the production of squeaks. Given that the PAG is thought to gate the production of all innate vocalizations,^1^ we hypothesized that courtship squeak-TRAPed PAG neurons would regulate the production of all squeaks, even those produced outside of a courtship context. We therefore also tested the effects of TRAP2-based PAG neuronal ablation on the production of footshock-elicited squeaks, as well as on the production of USVs (as a control vocalization type). Female TRAP2 mice received bilateral PAG injections of AAV-FLEX-caspase (day 0; targeted to ∼-4.0 mm A/P) and were single-housed on day 9, three days prior to the first behavioral test, to promote high rates of USV production during same-sex interactions (Fig. 5A).^17,18^ On day 12, single-housed females were given a same-sex interaction to elicit high rates of USVs, followed by mild footshocks (1.5s duration, 0.5 mA; see Methods) to measure footshock-elicited squeaks. Two days later, female subjects were given a 30-minute courtship interaction with a sexually experienced male to elicit high rates of courtship squeaks and then received an IP injection of 4-OHT to drive the ablation of courtship squeak-TRAPed PAG neurons. At least 10 days later, females were then re-tested in all three vocalization contexts. Control females were treated equivalently, except that they received bilateral PAG injections of AAV-FLEX-GFP. In pre-4-OHT behavioral measurements from TRAP-caspase and TRAP-GFP females, we found that the total number of courtship squeaks produced by each female was well related to total male mounting time (Fig. 5B), so male mounting time was used subsequently to normalize rates of female courtship squeaks in order to account for variability in male mounting times across trials (i.e., squeaks per second of male mounting). Although we did not measure or manipulate female estrous state, we found that normalized courtship squeak rates did not vary as a function of female rejection index (Fig. 5C; proportion of male mounting attempts rejected by the female, see Methods), consistent with past work showing that female mice produce squeaks across the estrous cycle during courtship interactions with males.^21,24^

**Figure 5.**
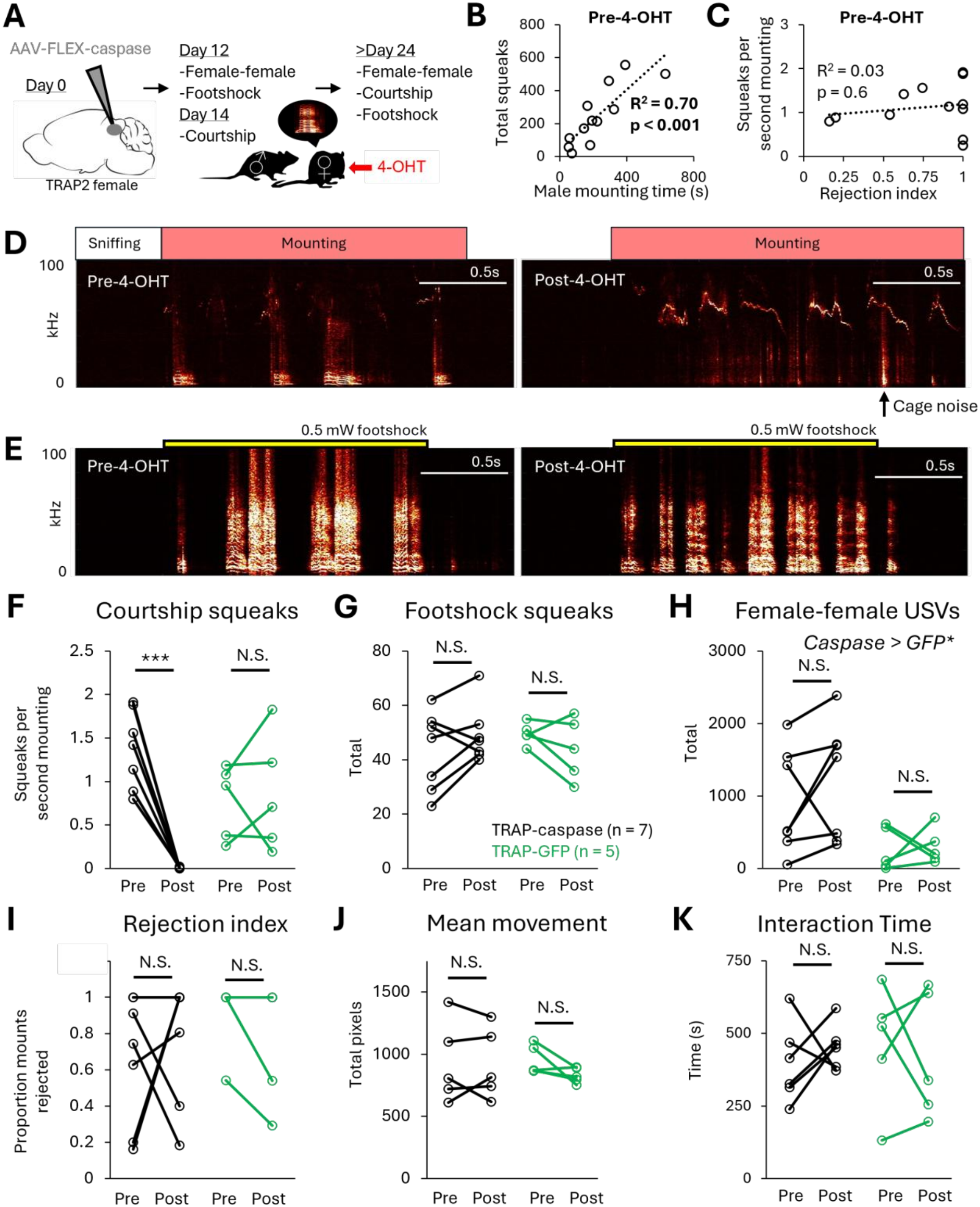
Caspase ablation of courtship squeak-TRAPed PAG neurons blocks the production of courtship USVs but does not affect rates of footshock-elicited squeaks or USVs. (A) Schematic shows experimental details and time course. (B) Plot shows the relationship between total male mounting time and total female courtship squeaks in pre-4-OHT courtship trials. (C) Plot shows the relationship of female reject index to normalized courtship squeak rates (squeaks per second male mounting) in pre-4-OHT courtship trials. (D) Left, representative spectrogram shows pre-4-OHT courtship squeaks produced by a TRAP-caspase female. Note that higher frequency USVs produced by the male are also visible in the spectrogram. Right, representative spectrogram shows the absence of female courtship squeaks following 4-OHT treatment in a TRAP-caspase female. (E) Same as D, for footshock-elicited squeaks. (F) Normalized courtship squeak rates are plotted for TRAP-caspase females (black symbols) and TRAP-GFP females (green symbols), pre– and post-4-OHT. (G) Same as (F), for total footshock-elicited squeaks. (H) Same as (F), for total female-female USVs. (I) Same as (F), for female rejection index. (J) Same as (F), for mean total footshock-elicited body movement. (K) Same as (F), for total time that subject females spent interacting with female visitor.

Following ablation of courtship squeak-TRAPed PAG neurons, TRAP-caspase females exhibited a striking reduction in the production of courtship squeaks, exhibiting near-zero rates following ablation (Fig. 5D,F; see also Videos S4-5). A two-way ANOVA with repeated measures on one factor revealed no significant main effect of group (F(1,10) = 0.38, p = 0.55), a significant main effect of time (F(1,10) = 18.62, p = 0.002) and a significant interaction between group and time (F(1,10) = 24.25, p < 0.001). Post-hoc tests showed that TRAP-caspase females exhibited a significant decrease in courtship squeak rates (Tukey’s HSD, p = 0.002), whereas courtship squeak rates did not change in TRAP-GFP females (p = 0.98). In contrast to courtship squeak production, female rejection index did not change for either group following 4-OHT treatment (Fig. 5I; F(1,10) = 0.00, p = 0.97 for main effect of group; F(1,10) = 0.53, p = 0.48 for main effect of time; F(1,10) = 3.95, p = 0.08 for interaction between group and time). In summary, ablation of courtship squeak TRAPed PAG neurons blocks the production of courtship squeaks but does not affect female non-vocal responses to male mounting attempts.

We hypothesized that ablation of courtship squeak-TRAPed PAG neurons would also block the production of squeaks elicited by mild footshock. Unexpectedly, ablation of courtship squeak-TRAPed PAG neurons had no effect on the total number of footshock-elicited squeaks (Fig. 5G; two-way ANOVA with repeated measures on one factor; F(1,10) = 0.01, p = 0.91 for main effect of group; F(1,10) = 0.00, p = 0.95 for main effect of time; F(1,10) = 3.94, p = 0.08 for interaction between group and time), nor was there a significant effect of ablation on the total amount of footshock-elicited body movement (see Methods; Fig. 5J; F(1,8) = 0.07, p = 0.80 for main effect of group; F(1,8) = 2.04, p = 0.19 for main effect of time; F(1,8) = 1.57, p = 0.24 for interaction between group and time; videos were not saved for n = 2 TRAP-caspase females, so footshock movement analysis includes only n = 5 TRAP-caspase females).

Similarly, ablation of courtship squeak-TRAPed PAG neurons had no significant effect on the total number of female-female USVs (Fig. 5H; two-way ANOVA with repeated measures on one factor; F(1,10) = 0.85, p = 0.38 for main effect of time; F(1,10) = 0.52, p = 0.49 for interaction between group and time), although pairs including TRAP-caspase females produced overall higher rates of USVs than pairs including TRAP-GFP females (F(1,10) = 6.30, p = 0.03 for main effect of group). Ablation of courtship squeak-TRAPed PAG neurons also had no significant effect on the total time that subject females spent investigating the visitor female (Fig. 5K; F(1,10) = 0.04, p = 0.86 for main effect of group; F(1,10) = 0.01, p = 0.91 for main effect of time; F(1,10) = 0.57, p = 0.47 for interaction between group and time; video was not saved for n = 1 TRAP-caspase female, so interaction time analysis includes only n = 6 TRAP-caspase females). In conclusion, ablation of courtship squeak-TRAPed PAG neurons blocks the production of courtship squeaks but has no effect on the total squeaks elicited by mild footshock or on USV production during same-sex interactions. These findings indicate that although courtship squeak-TRAPed PAG neurons critically regulate the production of female courtship squeaks, they are not required for squeak production across all behavioral contexts.

### Non-selective ablation of PAG neurons blocks the production of courtship squeaks and USVs but only attenuates the production of footshock-elicited squeaks

Given that the PAG is thought to gate the production of all innate vocalizations,^1–3^ we were surprised that ablation of courtship squeak-TRAPed neurons failed to block the production of footshock-elicited squeaks. We then conducted a final set of experiments to test whether the production of footshock-elicited squeaks depends on the PAG. Prior to receiving viral injections, 3-days single-housed female and male were given 30-minute interactions with a socially novel female to elicit high rates of USVs, female subjects only were given a 30-minute interaction with a sexually motivated male to elicit courtship squeaks, and both females and males were tested in a mild footshock paradigm to measure footshock-elicited squeaks (Fig. 6A). The next day, mice received bilateral PAG injections of a 1:1 mixture of AAV-Cre and AAV-FLEX-caspase viruses to drive non-selective ablation of PAG neurons (Fig. 6B). Viral injections were relatively large in volume (200 nL total), were targeted to approximately –4.0 mm A/P, and viral spread extended into the caudal-most portion of the PAG (Fig. S3). A minimum of 10 days later, subject mice were re-tested for all vocal behaviors.

**Figure 6.**
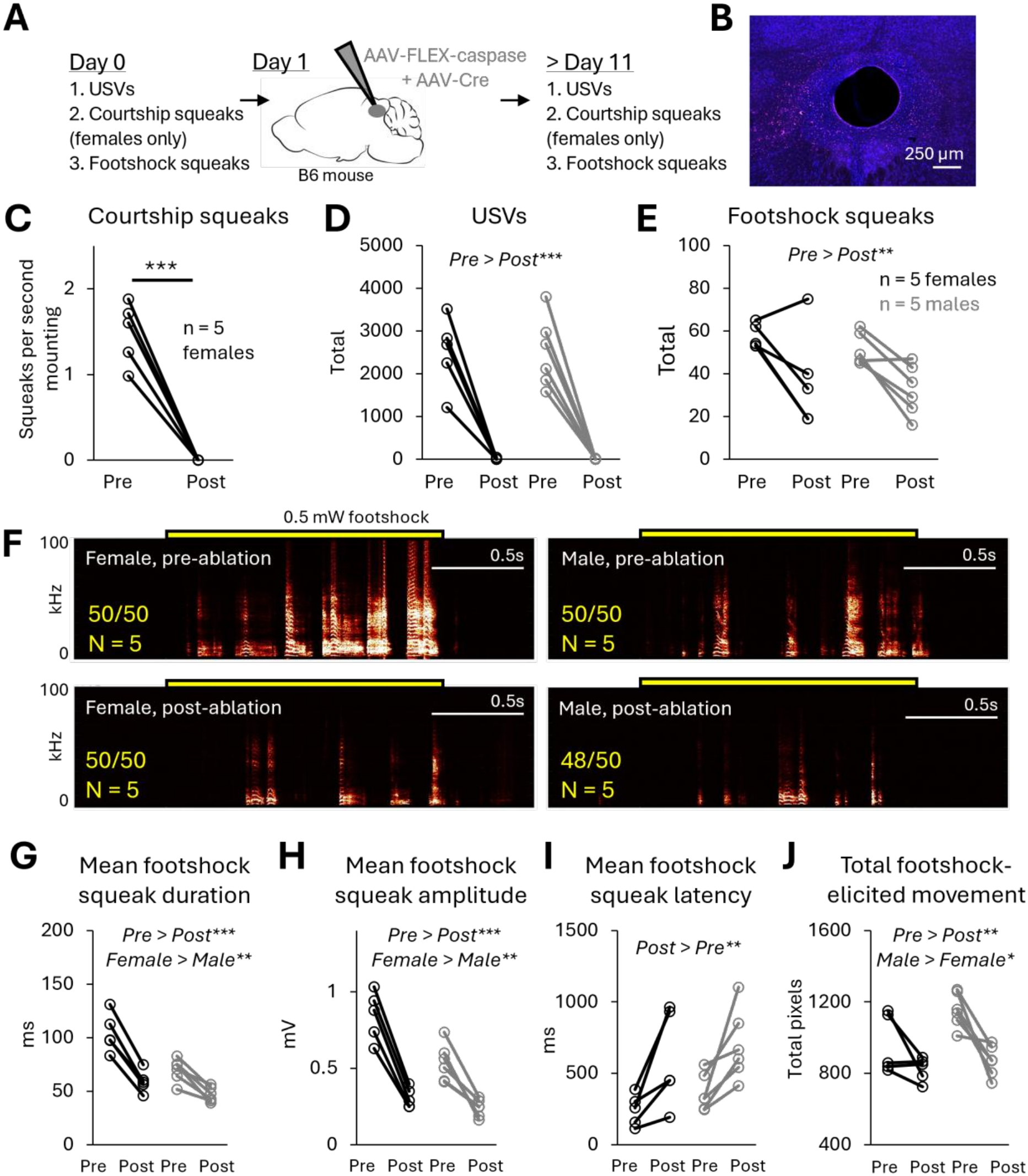
Non-selective ablation of PAG neurons blocks the production of courtship squeaks and USVs but only attenuates the production of footshock-elicited squeaks. (A) Schematic shows experimental details and time course. (B) Confocal image shows representative histology following non-selective ablation of PAG neurons. Note the presence of substantial cellular debris (seen as red autofluorescence) and enlarged ventricle. Blue, neurotrace. (C) Plot shows normalized courtship squeak rates produced by subject females pre– and post-ablation. (D) Plot shows total USVs produced by subject females (black symbols) and males (gray symbols) pre– and post-ablation. (E) Same as (D), for total footshock-elicited squeaks. (F) Representative spectrograms show footshock-elicited squeaks produced by a female (left column) and a male (right column) pre– and post-ablation. Yellow bars show timing of footshock. Yellow text indicates proportion of total trials across all mice in each group in which footshocks elicited squeaks. (G) Plots show mean duration of footshock-elicited squeaks produced by subject female and males pre– and post-ablation. (H) Same as (G), for amplitude of footshock-elicited squeaks. (I) Same as (G), for mean latency to footshock-elicited squeaks. (J) Same as (G), for mean total footshock-elicited body movement.

As expected, non-selective ablation of PAG neurons blocked the production of female courtship squeaks (Fig. 6C; paired t-test; t(4) = 10.38, p < 0.001) and blocked the production of USV in both female and male subjects (Fig. 6D; two-ANOVA; F(1,9) = 0.00, p = 1.00 for main effect of sex; F(1,9) = 96.94, p < 0.001 for main effect of time; F(1,9) = 0.00, p = 0.99 for interaction between sex and time). In contrast, PAG ablation attenuated but failed to block the production of footshock-elicited squeaks (Fig. 6E; F(1,9) = 0.25, p = 0.63 for main effect of sex; F(1,)) = 14.51, p = 0.004 for main effect of time; F(1,9) = 0.29, p = 0.60 for interaction between sex and time). The total number of footshock squeaks produced following non-selective PAG ablation was on average 63 ± 27% (mean ± stdev) of the number produced before ablation across all PAG-caspase mice. Aside from a decrease in the total number of footshock squeaks post-ablation, we also observed significant decreases in the mean duration and mean amplitude of footshock-elicited squeaks, as well as an increase in the mean latency from footshock onset to first squeak (Fig. 6G-I; two-way ANOVAs for each parameter; p ≤ 0.002 for main effect of time for all squeak features). In spite of the overall attenuation, mice continued to reliably produce squeaks in response to individual footshocks following non-selective PAG ablation (Fig. 6F; yellow text indicates proportion of total trials across all mice in each group in which footshocks elicited at least 1 squeak). In addition to changes in footshock-elicited vocal behavior, we found that the mean total footshock-elicited body movement was reduced following non-selective PAG neuronal ablation (Fig. 6K, Videos S6-7; F(1,9) = 19.10, p = 0.002 for main effect of time; F(1,9) = 7.29, p = 0.02 for main effect of sex; F(1,9) = 2.43, p = 0.15 for interaction between sex and time). Together, these data indicate that non-selective PAG ablation attenuates both vocal and non-vocal responses to footshock.

In summary, both vocal and non-vocal responses to footshock were significantly reduced following non-selective PAG ablation, but mice still continued to produce footshock-elicited squeaks. Taken together with the finding that PAG ablation abolished the production of courtship squeaks and USVs, these data support the idea that an intact PAG may not be required for the production of footshock-elicited squeaks (Fig. 7).

**Figure 7.**
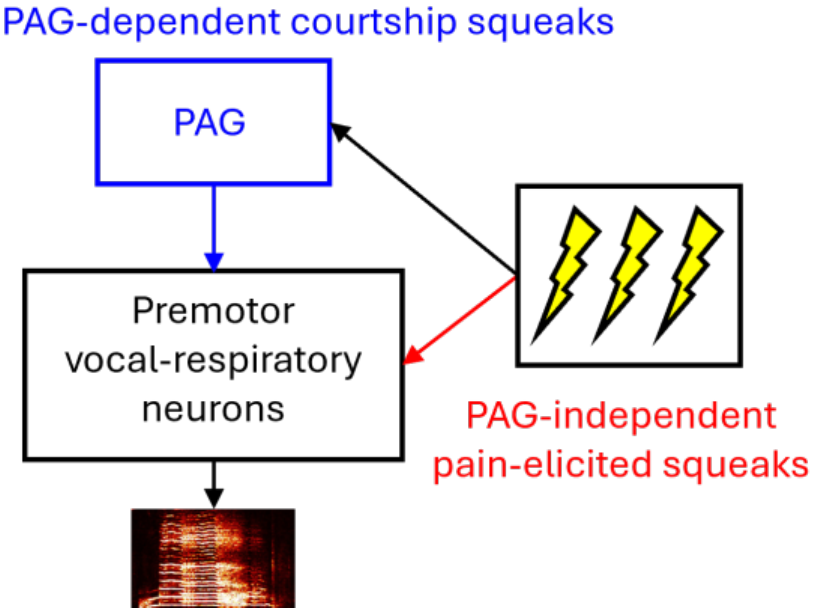
Model of PAG-dependent and PAG-independent pathways for squeak production. Activation of courtship squeak-TRAPed PAG neurons elicits squeaks, whereas ablation of courtship squeak-TRAPed PAG neurons blocks the production of courtship squeaks, indicating that a subset of PAG neurons critically regulates the production of female courtship squeaks (blue). In contrast, neither selective nor non-selective PAG neuronal ablation blocked the production of footshock-elicited squeaks, suggesting that a PAG-independent pathway may regulate the production of pain-elicited vocalizations (red).

## Discussion

In the current study, we used an activity-dependent labeling strategy to identify PAG neurons that critically regulate the production of courtship squeaks in female mice. Optogenetic activation of courtship squeak-TRAPed PAG neurons robustly elicits the production of squeaks, alongside variable non-vocal movements, and these TRAPed PAG neurons send axonal projections to hindbrain regions implicated in vocal production. In contrast, non-selective activation of glutamatergic PAG neurons weakly elicits the production of squeaks but robustly elicits escape-like locomotion in both females and males. Caspase-mediated ablation of courtship squeak-TRAPed PAG neurons selectively blocks the production of courtship squeaks but does not affect the production of USVs. Unexpectedly, ablation of courtship squeak-TRAPed PAG neurons failed to affect the production of footshock-elicited squeaks. Moreover, although non-selective ablation of PAG neurons blocked the production of both courtship squeaks in females and USVs in both sexes, this manipulation only attenuated but did not block the production of footshock-elicited squeaks. These experiments identify a population of PAG neurons that critically regulates the production of courtship squeaks in female mice, and our results indicate that the production of pain-elicited vocalizations may be regulated by a non-canonical, PAG-independent pathway.

Optogenetic activation of courtship squeak-TRAPed PAG neurons might promote squeak production via recruitment of pontine and/or hindbrain vocalization-related neurons (Fig. 3). Of particular interest are the axonal projections of these neurons to RAm, which has been implicated in regulating the production of both USVs and squeaks,^30,32,35–37^ as well as axonal projections to the peri-LC region, which has been implicated in regulating USV production,^34^ although the role of this region in regulating squeak production remains untested. Alternatively, activation of TRAPed PAG neurons might promote the production of squeaks in a less direct manner by driving changes in female internal state. Our experiments provide mixed support for these ideas. On the one hand, the finding that optogenetic activation of TRAPed PAG neurons elicits robust squeaks but variable effects on movement across females and also fails to elicit male-directed rejection behaviors seems inconsistent with the idea that optogenetically-elicited squeaks are produced secondarily to a change in female internal state. On the other hand, we observed freezing behavior following optogenetic activation of TRAPed PAG neurons that persisted in between optogenetic stimulations. This result suggests that optogenetic activation exerts at least some effects on internal state, although it remains unclear whether these effects are a direct result of PAG neuronal activation vs. a secondary consequence of optogenetically-elicited behaviors on internal state (e.g., producing a squeak unexpectedly may scare the mouse). Another issue that warrants future study is whether the effects of optogenetic activation on vocal and non-vocal behaviors are driven by a single population of TRAPed PAG neurons that influence multiple aspects of behavior, or alternatively, whether courtship squeak-TRAPed PAG neurons represent multiple populations with distinct effects on behavior. Given that non-selective activation of VGlut2^+^ PAG neurons drives robust locomotor effects and also drives freezing in between trials (Fig. 4F) and given also past work showing that activation of the dorsal half of the PAG can be used as an unconditioned stimulus in fear-conditioning paradigms,^40–42^ it is likely that even a small amount of off-target viral labeling in our experiments could drive the locomotor effects that we observed in our TRAP-ChR2 females. Future experiments with single-cell resolution of PAG neural activity will be important to test whether there exist PAG neurons with motoric-like, squeak-related activity. Future work can also test whether activation of projection-defined subsets of courtship squeak-TRAPed neurons, or their axon terminals in downstream pontine and hindbrain regions, is sufficient to selectively elicit the production of squeaks.

Ablation of courtship squeak-TRAPed PAG neurons failed to affect rates of female USV production (Fig. 5H). Together with recent work showing that ablation of PAG-USV neurons does not block the production of squeaks,^23^ this finding adds further support to the idea that distinct populations of PAG neurons regulate the production of USVs and squeaks. Given that USVs are produced via an intralaryngeal whistle mechanism^43–45^ and squeaks are produced via vibration of the vocal folds, one idea is that non-overlapping populations of PAG neurons regulate the production of vocalization types that differ in their phonation mechanism. A recent study in the singing mouse (*Scotinymus teguina*) found that ablation of caudolateral glutamatergic PAG neurons blocked the production of both USVs and territorial songs, both of which are produce via intralaryngeal whistle.^46,47^ Although it remains to be determined whether USV-related and song-related PAG neurons represent overlapping populations vs. distinct but spatially-intermingled populations, these results raise the related possibility that shared populations of PAG neurons may regulate vocalization types that share a phonation mechanism.

Nonetheless, in contrast to PAG-USV neurons, which critically regulate the production of USVs across multiple behavioral contexts,^23,25^ the PAG neurons characterized in the current study do not represent all-purpose squeak-promoting neurons, because ablation of courtship squeak-TRAPed PAG neurons did not affect the production of footshock-elicited squeaks (Fig. 5G). Do courtship squeak-TRAPed PAG neurons selectively regulate the production of female courtship squeaks, or might they regulate squeaks produced in additional behavioral contexts? A recent study found the pharmacological silencing of the dorsolateral PAG blocked the production of squeaks elicited by tail suspension in mice bred for high-anxiety behaviors.^22^ Thus, one idea is that courtship squeak-TRAPed PAG neurons may critically regulate all squeaks produced in non-pain-associated behavioral contexts, although testing this idea would require the identification of additional, non-pain-associated contexts that reliably elicit the production of squeaks in B6 females. In a similar vein, although we used the courtship squeak paradigm to identify squeak-associated PAG neurons, we do not think that the identified neurons are female-specific. In line with this idea, optogenetic activation of VGlut2^+^ PAG neurons elicited the production of squeaks in both females and males, albeit much more weakly than TRAP-based optogenetic activation (Fig. 4). Future work to further characterize courtship squeak-TRAPed PAG neurons in females would set the stage to test whether similar neurons exist in the PAG of male mice, and if so, whether those neurons regulate squeak production in males.

Given that the PAG is thought to regulate the production of all innate vocalizations in mammals,^1^ the finding that even non-selective PAG neuronal ablation fails to block the production of footshock-elicited squeaks was unexpected. These results suggest that a non-canonical, PAG-independent pathway may regulate the production of footshock-elicited squeaks, and perhaps pain-elicited vocalizations more generally. Recent work has implicated the hindbrain region nucleus retroambiguus (RAm) in regulating the production of squeaks,^30,37^ including the finding that blocking neurotransmitter release from RAm neurons can block the production of squeaks elicited by tail pinch.^32^ One possibility is that ascending pain information might access RAm independently of the PAG (Fig. 7), either directly via spinal inputs to the reticular formation^48–51^ or indirectly through an intermediate region such as the parabrachial nucleus, which is important for pain processing^28,29,51^ and provides input to RAm.^52^ Future work identifying neuronal populations that provide input to squeak-related RAm neurons, as well as identifying additional hindbrain premotor neurons that regulate squeak production, could shed light on circuits that may promote the production of pain-elicited vocalizations in a PAG-independent manner.

Most studies to-date investigating brainstem vocalization circuits in rodents have focused on neurons that regulate USV production.^23,25,32,35–37,39,53–55^ However, there may be important differences in the neural circuits that regulate the production of ultrasonic vs. voiced vocalizations, and the neural circuits that regulate the production of squeaks are only beginning to be explored. The identification of PAG neurons that regulate the production of female courtship squeaks provides an entry point to a detailed characterization of the brainstem circuits that regulate squeak production, which will in turn contribute to our understanding of how brainstem vocalization circuits enable the production of different vocalization types.

## Acknowledgements

This work was funded by NIMH (R01MH136887). We thank the CARE staff for their excellent mouse husbandry.

## Author contributions

DJZ, RH, and PZ conducted experiments and analyzed data. ZZ analyzed data. KAT designed the experiments, analyzed data, and wrote the paper, and all authors approved of the final version.

## Supplemental Information

**Figure S1. Comparison of optogenetically-elicited movements across experimental groups.** (A) Plots show trajectories of optogenetically-elicited movements for the n=5 TRAP-ChR2 females that exhibited optogenetically-elicited squeaks. (B) Same as (A), for the n = 3 TRAP-ChR2 females that did not exhibit optogenetically-elicited squeaks. (C) Same as (A), for TRAP-GFP control females. (D) Same as (A), for the n = 7 VGlut2-ChR2 mice that exhibited optogenetically-elicited squeaks.

**Figure S2. Comparison of optogenetically-elicited squeaks and body movements in solo vs. male optogenetic sessions.** (A) Rates of optogenetically-elicited squeaks are plotted for TRAP-ChR2 females across solo and male sessions. (B) Same as (A), for optogenetically-elicited movement. (C) Plots show trajectories of optogenetically-elicited movements for each TRAP-ChR2 female in the solo session (top row) and the male session (bottom row).

**Figure S3. Confocal images of PAG sections following non-selective PAG neuronal ablation.** (Representative confocal images show the PAG following non-selective neuronal ablation. Blue, neurotrace. Red, autofluorescence. Numbers on images indicate position relative to bregma. Scale bars, 250 µm.

**Video S1. Optogenetic activation of courtship squeak-TRAPed PAG neurons elicits the production of squeaks.** Video shows an example optogenetic stimulation (1s-long, 10 mW, 33 Hz) delivered to a solo-tested TRAP-ChR2 female.

**Video S2. Optogenetic activation of courtship squeak-TRAPed PAG neurons does not elicit male-directed rejection behaviors.** Video shows an example optogenetic stimulation (1s-long, 10 mW, 33 Hz) delivered to a TRAP-ChR2 female tested in the presence of a male. Same TRAP-ChR2 female is shown in Video S2 and S1.

**Video S3. Optogenetic activation of VGlut2^+^ lateral PAG neurons weakly elicits squeaks and robustly elicits escape-like locomotion.** Video shows an example optogenetic stimulation (3s-long, 5 mW, 33 Hz) delivered to a solo-tested VGlut2-ChR2 male.

**Video S4. Production of courtship squeaks by a TRAP-caspase female, prior to ablation of courtship squeak-TRAPed PAG neurons.** Video shows the production of courtship squeaks by a TRAP-caspase female, following viral injection but prior to treatment with 4-hydroxytamoxifen.

**Video S5. Ablation of courtship squeak-TRAPed PAG neurons abolishes the production of courtship squeaks.** Video shows the absence of courtship squeaks during an interaction with a sexually motivated female. Same TRAP-caspase female is shown as in Video S4, post-4-hydroxytamoxifen treatment.

**Video S6. Production of footshock squeaks prior to non-selective ablation of PAG neurons.** Video shows footshock squeaks produced by a female mouse prior to non-selective ablation of PAG neurons.

**Video S7. Production of footshock squeaks following non-selective ablation of PAG neurons.** Video shows footshock squeaks produced by a female mouse following non-selective ablation of PAG neurons. Same mouse is shown in Videos S6 and S7.

## Methods

### Resource Availability

#### Lead contact

Further information and requests for resources should be directed to and will be fulfilled by the lead contact, Dr. Katherine Tschida.

#### Material availability

No new materials were generated for this study.

#### Data and code availability

All source data and Matlab codes used to analyze data are available upon request.

##### Experimental model and study participant details

#### Ethics statement

All experiments and procedures were conducted according to protocols approved by the Cornell University Institutional Animal Care and Use Committee (protocol #2020-001).

#### Subject details

The following mouse strains were used in this study: C57BL/6J females and males (Jackson laboratories, 000664), for the PAG Fos experiments in Fig. 1 (females only) and for the PAG Cre-caspase experiments in Fig. 6 (both sexes); TRAP2 females (Jackson Laboratories, 030323), for the TRAP-ChR2 and TRAP-caspase experiments in Figs. 2, 4, and 5; VGlut2-Cre females and males (Jackson Laboratories, 028863), for the optogenetic activation experiments in Fig. 3. All subject mice were housed with their siblings and both parents until weaning at postnatal day 21. Mice were kept on a 12:12 reversed light/dark cycle, were housed in ventilated micro-isolator cages in a room with regulated temperature and humidity and were provided with unrestricted access to food and water. A running wheel (Innovive) was present in all home cages from the time of weaning and was subsequently removed immediately prior to any behavioral trials that were conducted in the subject mouse’s home cage. All mice were at least 8 weeks old at the time of surgeries and/or experiments. All female mice used in these experiments were naturally cycling, and estrous state was not tracked.

### Method details

#### Courtship squeak trials

To elicit courtship squeaks, 3-days-single-housed female subject mice were placed in the home cage of a sexually experienced male for a courtship trial (1-30 minutes in duration). Male home cages were placed in a sound-attenuating chamber (Med Associates) equipped with an ultrasonic microphone (Avisoft, CMPA/CM16), an infrared light source (Tendelux), and a webcam (Logitech, with the infrared filter removed to enable video recording under infrared lighting conditions). The cage lid was removed, and the cage was fitted with a custom lid that allowed microphone access and permitted audio and video recordings. All materials within the cage were removed, including bedding, to reduce low-frequency noise and facilitate the detection of female courtship squeaks. Audio recordings were amplified (Presonus TubePreV2) and digitized at 250 kHz (Spike 7, CED).

#### USV trials

To elicit USVs from female or male subject mice, 3-days-single-housed subject mice were given a 30-minute social interaction with a socially novel, group-housed female in their home cage. Home cages were placed in a sound-attenuating chamber (Med Associates) and fitted with custom lids that allowed microphone access and permitted audio and video recordings. Running wheels were removed but bedding was left in place. Audio recordings were made using an ultrasonic microphone (Avisoft, CMPA/CM16), amplified (Presonus TubePreV2), and digitized at 250 kHz (Spike 7, CED). Video recordings were made using a webcam (Logitech) that was modified by removing the infrared filter to permit video recordings under infrared lighting conditions.

#### Footshock trials

Subject mice were placed in a footshock chamber (Med Associates) inside a sound attenuating chamber (Med Associates) equipped with an ultrasonic microphone (Avisoft, CMPA/CM16), an infrared light source (Tendelux), and a webcam (Logitech, with the infrared filter removed to enable video recording under infrared lighting conditions). A mild (0.5 mA, 1.5s-long) shock was delivered once every 30 seconds, for a total of 10 footshocks (trial duration = 5 minutes).

#### Detection of vocalizations from audio recordings

In footshock and courtship trials (females only), a custom Matlab code was used that allowed trained users to manually annotate the onsets and offsets of individuals squeaks from spectrograms created from audio recordings. USVs were detected with custom Matlab codes (Tschida et al., 2019) using the following parameters: mean frequency > 45 kHz; spectral purity > 0.3; spectral discontinuity < 1.0; minimum USV duration = 5 ms; minimum inter-syllable interval = 30 ms).

#### Viruses

The following viruses were used: AAV-2/1-EF1alpha-FLEX-hChR2(h134R)-EYFP-WPRE-HGHp (abbreviated as AAV-FLEX-ChR2; Addgene #20298); AAAV-2/1-CAG-FLEX-EGFP-WPRE (AAV-FLEX-GFP; Addgene #51502); AAV-2/1-CAG-GFP (AAV-GFP; Addgene #37825); AAV-2/5-EF1alpha-FLEX-taCasp3-TEVp (AAV-FLEX-caspase; Addgene #45580); AAV-2/1-hSyn-Cre.WPRE.hgGH (AAV-Cre; Addgene #105553).

Subject females in the TRAP2-ChR2 experiments received unilateral 100 nL PAG injections of AAV-FLEX-ChR2 (experimental females) or AAV-FLEX-GFP (control females). Subject males and females in the VGlut2-ChR2 experiments received unilateral 50 nL PAG injections of AAV-FLEX-ChR2 (experimental mice) or AAV-FLEX-GFP (control mice). Subject females in the TRAP-caspase experiments received bilateral 200 nL PAG injections of a mixture (10:1) of AAV-FLEX-caspase (to ablate recently active PAG neurons) and AAV-GFP (to visualize injection targeting post-hoc). Control females in the TRAP-caspase experiments received bilateral 200 nL PAG injections of AAV-FLEX-GFP. Subject females and males in the Cre-caspase experiment received bilateral PAG injections of 200 nL of a mixture (1:1) of AAV-FLEX-caspase and AAV-Cre.

#### Stereotaxic surgery

Mice were anesthetized using isoflurane (2.5% for induction, then 1.5 – 2.5% for maintenance) and then securely positioned in a stereotaxic apparatus (Angle Two, Leica). A midline incision in the scalp was made to expose the skull, and small craniotomies were created dorsal to each injection site using a surgical drill. The PAG was targeted by making a craniotomy rostral to the intended injection coordinates and lowering the glass pipette containing virus into the brain, angled caudally at an angle of 13-25 degrees relative to vertical. Viruses were pressure-injected with a Nanoject III (Drummond) at a rate of 5 nL every 20 seconds. Viral injection pipettes were left in place for a minimum of 5 minutes before and after viral injections to minimize backflow upon pipette withdrawal from the brain. Surgical sutures (MV-711-V, Oasis) and tissue adhesive (3M) were used to close the incision.

For optogenetic activation experiments, an optogenetic ferrule (RWD Fiber Optic Cannula, Ø1.25 mm Ceramic Ferrule, 200 µm Core, 0.22 NA, length = 3 mm) was implanted above the viral injection site immediately following the viral injection and was secured in place with Metabond (Parkell).

#### TRAP activity-dependent labeling

4-hydroxytamoxifen (4-OHT, HelloBio, HB6040) was dissolved at 20 mg/mL in ethanol by shaking at 37°C and was then aliquoted (75 uL) and stored at –20°C. Before use, 4-OHT was redissolved in ethanol by shaking at 37°C and filtered corn oil was added (Sigma, C8267, 150 uL). Ethanol was then evaporated by vacuum under centrifugation to give a final concentration of 10 mg/mL, and 4-OHT solution was used on the same day it was prepared, delivered at a final dosage of 50 mg/kg.

To express viral transgenes in recently active neurons, we used the Targeted Recombination in Active Populations (TRAP2) strategy. Two weeks following viral injection, TRAP2 females were single-housed for 3 days and then given 30-minute courtship trials to elicit high rates of courtship squeaks (as described above). Following the courtship trial, females received I.P. injections of 4-OHT (150 uL of 10 mg/mL 4-OHT in filtered corn oil) to enable expression of viral transgenes in recently active PAG neurons. Females remained single-housed following the TRAPing session and until all behavioral tests were completed.

#### Optogenetic activation

Subject mice were placed alone in a clean testing chamber for a 5-minute habituation period after connecting the laser patch cable to the female’s optogenetic ferrule. PAG neurons were optogenetically activated via a 473 nm laser (5-10 mW, 20-33 Hz, 1-3s-long trains) approximately once every 30 seconds. Laser stimuli were driven by computer-controlled voltage pulses (Spike 2). Optogenetic activation trials began 2 weeks following 4-OHT treatment (a total of 4 weeks following viral injection) for TRAP-ChR2 females and 3 weeks following viral injection for VGlut2-Cre mice.

#### Neuronal ablation

To test the effects of selective (TRAP2-based) and non-selective PAG neuronal ablation of vocal production, subject mice were given courtship trials to elicit courtship squeaks (female subjects only), interactions with group-housed females to elicit USV production, and footshock trials to elicit footshock squeaks. Total vocalizations of each type for each subject mouse was compared pre-ablation and post-ablation to assess the effects of PAG neuronal ablation. Post-ablation tests were conducted 10 days following 4-OHT treatment (∼3 weeks following viral injection) for TRAP-caspase females and a minimum of 10 days following viral injection for Cre-caspase females and males.

#### Fos immunohistochemistry

Two hours following the start of behavior trial, subject mice were deeply anesthetized using isoflurane and then transcardially perfused with phosphate-buffered saline (PBS, pH 7.4), followed by 4% paraformaldehyde (PFA, in 0.1 M PBS, pH 7.4). Brains were subsequently dissected and post-fixed in 4% PFA for 24 hours at 4°C, followed by immersion in 30% sucrose solution in PBS for 48 hours at 4°C. Afterward, brains were embedded in frozen section embedding medium (Surgipath, VWR), flash frozen in a dry ice-ethanol (100%) bath, and then stored at ^-^80°C until sectioning. Coronal sections were cut on a cryostat to a thickness of 80 μm, washed in PBS (3 x 5 mins at RT), permeabilized for 2-3 hours in PBS containing 1% Triton X-100 (PBST), and then blocked in 0.3% PBST containing 10% Blocking One (Nacalai USA) for 1 hour at RT on a shaker. Sections were then incubated for 24 hours at 4°C with primary antibody in blocking solution (1:1000 rabbit anti-Fos, Cell Signaling Technologies, 2250S), washed 3 x 10 minutes in 0.3% PBST, then incubated for 24 hours at 4°C with secondary antibody in blocking solution (1:1000, Alexa Fluor 488 or Alexa Fluor 633 goat-anti-rabbit, Invitrogen, plus 1:500 NeuroTrace, Invitrogen). Finally, sections were washed for 2 x 10 minutes in 0.3% PBST, followed by washing for 2 x 10 minutes in PBS. After mounting on slides, sections were dried and coverslipped with Fluromount G (Southern Biotech). Slides were imaged with a 10x objective on a Zeiss 900 laser scanning confocal microscope.

Quantification of courtship-squeak associated Fos expression was performed on coronal sections of the PAG, spanning –3.4 mm to –5 mm caudal to bregma. A trained scorer used ImageJ to draw a horizontal line that demarcated the dorsal two-thirds of the PAG in each section (ventral third of PAG not scored). Within the dorsal two-thirds of the PAG, Fos-positive neurons were then manually annotated using ImageJ.

#### Axon projection quantification

This analysis included midbrain and hindbrain coronal sections from n = 5 TRAP-ChR2 females in which optogenetic activation of courtship squeak-TRAPed PAG neurons elicited opto-squeaks. Brains were processed as detailed above, and 80 μm-thick coronal sections were imaged with a 10x objective on a Zeiss 900 laser scanning confocal microscope. First, brain regions of interest (ROIs) were manually annotated in a representative plane of section using ImageJ, both ipsilateral and contralateral to the side of the PAG viral injection. To detect axons in each ROI, images were binarized using ImageJ’s threshold tool to accept pixels whose values were equal to or greater than the value at the right-most edge of the histogram visualizing the distribution of pixel intensities for that section. The measurement tool was then used to calculate the area and the integrated pixel density of each ROI. Dividing the integrated pixel density by 255 yields the total number of pixels in each ROI that passed threshold, and the axon projection density was calculated for each ROI by dividing this number of pixels by the total area of the ROI. The following ROIs were quantified: (1) pontine reticular formation, (2) lateral parabrachial nucleus, (3) peri-locus coeruleus region, (4) lateral paragigantocellular nucleus (around the level of 7N), (5) gigantocellular reticular nucleus, alpha part (around the level of 7N), (6) gigantocellular reticular nucleus, ventral part (around the level of nucleus ambiguus), and (7) nucleus retroambiguus.

#### Quantification of acoustic features of optogenetically-elicited squeaks and naturally-produced courtship squeaks

We used custom Matlab codes to generate spectrograms of our audio recordings and manually annotate the onset and offset of each squeak. For all annotated squeaks, we calculated 3 acoustic features: (1) duration, (2) dominant frequency (calculated at each time point of the squeak and then averaged across the squeak), and (3) amplitude. Amplitude is reported in volts, directly measured as the mean amplitude of the audio waveform recorded in Spike2 during manually annotated squeak times. All such comparisons used audio recordings conducted in the same behavioral chamber and with the microphone set in the same position across trials.

#### Analyses of non-vocal social behaviors

Trained scorers used BORIS software (v.8.13; Friard and Gamba, 2016) to score the following non-vocal behaviors: (1) male mounting (in courtship trials, used to normalize total number of female courtship squeaks across trials) and (2) subject-initiated social investigation of a novel, group-housed female (in USV trials). Social investigation included sniffing and following. Mounting was defined as the male mouse positioning its forelimbs on top of the body of the female, sometimes with pelvic thrusts and sometimes without.

In courtship trials for TRAP-caspase and TRAP-GFP females, female rejection index was calculated as the proportion of total male mounting attempts that were rejected by the female in each trial. A trained observer watched portions of the video previously scored as containing male mounting attempts. The outcome of each attempt was then scored as either female rejection (defined as the female fleeing from the male, exhibiting active rejection behaviors such as kicking or slapping, re-orienting her body to avoid the mounting attempt, and/or jumping on top of the cage air inlet port to terminate the mounting attempt) or female acceptance (defined as the female remaining in place as the male attempted to mount and/or receiving an intromission).

In some trials, total movement (either footshock-elicited or optogenetically-elicited) was estimated using a custom MATLAB code that allows the user to mark the position of a mouse (marked as either the position of the base of the optogenetic implant or the top of the head midway between the ears) in every 6^th^ frame (i.e., once per 200 ms). Total movement was then calculated as the sum of changes in position (in pixels) across pairs of marked frames that followed the onset of the footshock or laser stimulation.

### Quantification and statistical analysis

The Shapiro-Wilk test was used to evaluate the normality of data distributions. Parametric, two-sided statistical comparisons were used in all analyses of normally distributed data, and non-parametric, two-sided comparisons were used in analyses that included non-normally distributed data. In all cases, the significance level was set as alpha = 0.05. No statistical methods were used to pre-determine sample size. Errors represent standard error of the mean unless otherwise noted. Statistical analyses were carried out using R 4.4.1 (R Core Team, 2024) and R Studio 2026.01.1 +403 (Posit team, 2026). Same-sex cages of mice were selected at random for inclusion into experimental groups. Mice were excluded from the study only if their viral injections were mis-targeted.

