## Supplemental figures for "Midbrain neurons regulate the production of courtship squeaks in female mice"

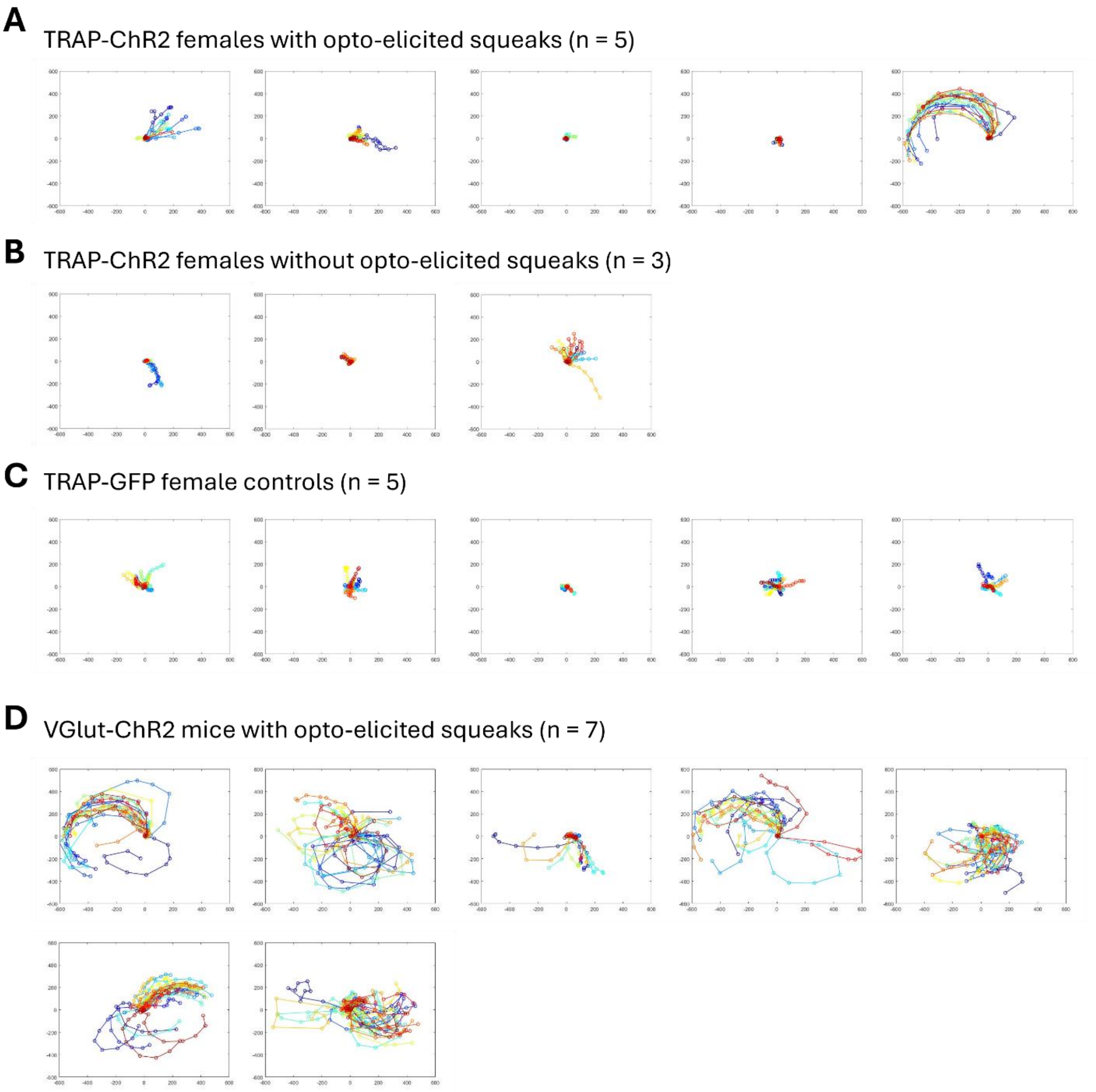


**Figure S1. Comparison of optogenetically-elicited movements across experimental groups.** (A) Plots show trajectories of optogenetically-elicited movements for the n = 5 TRAP-ChR2 females that exhibited optogenetically-elicited squeaks. (B) Same as (A), for the n = 3 TRAP-ChR2 females that did not exhibit optogenetically-elicited squeaks. (C) Same as (A), for TRAP-GFP control females. (D) Same as (A), for the n = 7 VGlut2-ChR2 mice that exhibited optogenetically-elicited squeaks.

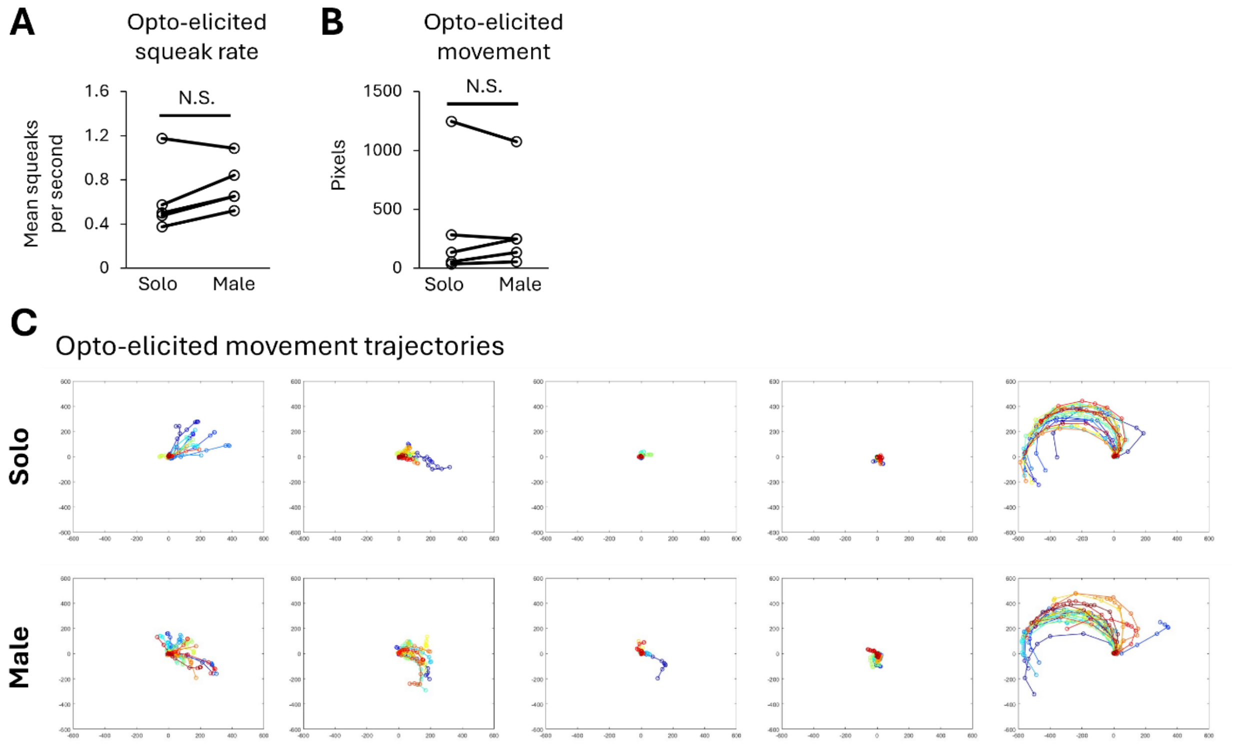


**Figure S2. Comparison of optogenetically-elicited squeaks and body movements in solo vs. male optogenetic sessions**. (A) Rates of optogenetically-elicited squeaks are plotted for TRAP-ChR2 females across solo and male sessions. (B) Same as (A), for optogenetically-elicited movement. (C) Plots show trajectories of optogenetically-elicited movements for each TRAP-ChR2 female in the solo session (top row) and the male session (bottom row).

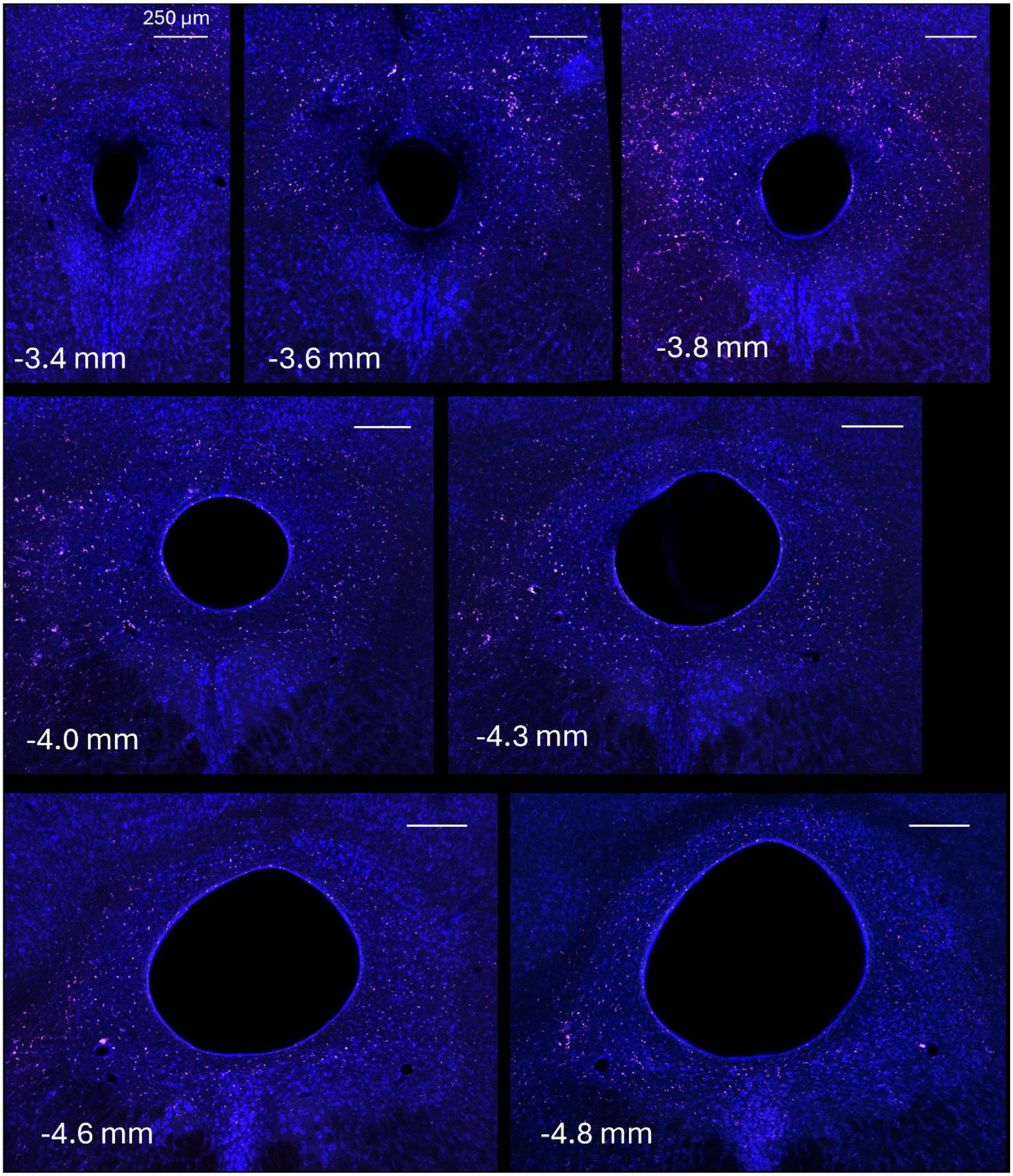


**Figure S3. Confocal images of PAG sections following non-selective PAG neuronal ablation**. Representative confocal images show the PAG following non-selective neuronal ablation. Blue, neurotrace. Red, auto-fluorescence. Numbers on images indicate position relative to bregma. Scale bars, 250 µm.
